# Reconstruction of resting-state networks from macaque electrocorticographic data

**DOI:** 10.1101/221051

**Authors:** R. Hindriks, C. Micheli, C.A. Bosman, R. Oostenveld, C. Lewis, D. Mantini, P. Fries, G. Deco

## Abstract

The discovery of haemodynamic (BOLD-fMRI) resting-state networks (RSNs) has brought about a fundamental shift in our thinking about the role of intrinsic brain activity. The electrophysiological underpinnings of RSNs remain largely elusive and it has been shown only recently that electrophysiological cortical rhythms are organized into RSNs. Most electrophysiological studies into RSNs use magnetoencephalography (MEG) or electroencephalography (EEG), which limits the spatial scale on which RSNs can be investigated. Due to their close proximity to the cortical surface, electroencephalographic (ECoG) recordings can potentially provide a more detailed picture of the functional organization of resting-state cortical rhythms. In this study we propose using source-space independent component analysis for identifying generators of resting-state cortical rhythms as recorded with ECoG and reconstructing their network structure. Their network structure is characterized by two kinds of connectivity: instantaneous correlations between band-limited amplitude envelopes and oscillatory phase-locking. Using simulated data, we find that the reconstruction of oscillatory phase-locking is more challenging than that of amplitude correlations, particularly for low signal-to-noise levels. Specifically, phase-lags can both be over- and underestimated as a consequence of first-order and higher-order volume-conduction effects, which troubles the interpretation of interaction measures based on imaginary phase-locking or coherence. The methodology is applied to resting-state beta (15-30 Hz) rhythms within the motor system of a macaque monkey and leads to the identification of a functional network of seven cortical generators that are distributed across the sensorimotor system. The spatial extent of the identified generators, together with consistent phase-lags, suggests that these rhythms can be viewed as being spatially continuous with complex dynamics including traveling waves. Our findings illustrate the level of spatial detail attainable with source-projected ECoG and motivates wider use of the methodology for studying resting-state as well as event-related cortical dynamics in macaque and human.

## 1 Introduction

Functional magnetic resonance imaging (fMRI) has demonstrated that in the absence of structured stimuli and explicit cognitive tasks i.e. during the resting-state, hemodynamic fluctuations in cortical gray-matter of humans and macaques are correlated across regions and constitute resting-state networks (RSNs) [20, 8, 14, 36]. The electrophysiological counterparts of hemodynamic RSNs have only recently been discovered using scalp electroencephalography (EEG) and magnetoencephalography (MEG) [38, 9, 2, 28, 34]. The high temporal resolution of electrophysiological recording techniques and the rich dynamics of such data allow for investigating the spectral signatures of hemodynamic RSNs. The spatial resolution of scalp EEG and MEG is limited, however, due to the large distance of the sensors to the cortex (in case of MEG) and the spatial lowpass filtering properties of the skull (in case of scalp EEG) [22, 41]. Electrocorticography (ECoG), which refers to voltage measurements taken directly from the cortical surface, have higher spatial resolution than scalp EEG and therefore allow studying resting-state cortical activity in high temporal and spatial resolution [25, 21]. ECoG measurements are often analyzed on the sensor-level and is it commonly believed that source-modeling does not improve the spatial resolution, because the electrodes are near the cortical surface. The human cortex - and to a lesser extent, macaque cortex - is convoluted with sulci that can be up to three centimeters deep, which complicates the interpretation of sensor-level ECoG measurements. This motivates the development of volume conductor models and source-modeling methods for ECoG measurements and it has been shown that source-modeling of ECoG measurements is feasible and does in fact improve spatial resolution [17, 56, 6, 33, 11, 47, 7].

The potential of source-modeling of ECoG measurements has only began to be explored. In particular, existing studies have exclusively focused on epileptic cortical activity in human subjects [17, 56, 6, 33, 11, 47, 7]. In this study, we investigate the possibility of reconstructing resting-state networks from macaque ECoG measurements. If feasible, this opens up new perspectives for studying the electrophysiological underpinnings of hemodynamic RSNs with higher spatial resolution than offered by scalp EEG and MEG. Within the EEG/MEG resting-state literature, two main methods are used to reconstruct RSNs from source-modeled data: seed-based correlation analysis [28, 42] and decomposition approaches, most notably independent component analysis (ICA) [37, 1, 32, 34]. In this study, we exclusively focus on spatial ICA. EEG and MEG electrophysiological resting-state studies commonly focus on correlations between band-limited amplitudes of source-projected data, as these have been shown to constitute RSNs. Besides amplitude correlations, we also assess the feasibility of reconstructing oscillatory phase-locking, as the latter maximally exploits the temporal richness of electrophysiological recordings and has been hypothesized to constitute a basic mechanism of cortical communication [15, 16]. We simulate networks of oscillatory generators and assess the effect of different signal-to-noise ratios and of the selected number of independent components. We illustrate the methodology on ECoG measurements of a macaque monkey displaying strong sensorimotor beta (15 – 30 Hz) oscillations.

## 2 Materials and Methods

### 2.1 Recordings and pre-processing

Data were recorded on two different resting-state sessions from a macaque monkey using a 252-electrode subdural electrode grid covering a large part of the left hemisphere [49]. During the two recording sessions, which we will refer to as session 1 and session 2, the monkey was situated in front of a wide screen in an otherwise dark room. Simultaneously recorded electro-ocular signals were inspected for the presence of eye-blinks and saccades, and confirmed that the monkey was awake with eyes open throughout both sessions. The duration of session 1 and 2 was 11.5 and 13 minutes, respectively. Because a substantial number of electrodes overlying posterior parts of the cortex were noisy and devoid of oscillatory activity, analysis was confined to the 122 most anterior channels. Remaining noisy electrodes (two and one in session 1 and 2, respectively) were replaced by their neighbor averages. The signals were filtered using a notch filter at 50 Hz, a fourth-order zero-phase Butterworth lowpass filter with cut-off frequency 60 Hz, and subsequently down-sampled to 500 Hz and converted to average montage. We selected five consecutive minutes of artifact-free data of each session. The electrode-averaged power spectra of the selected data are shown in Figure 1A. Figure 1B shows ten-second time-series from the 20 most anterior electrodes. In both sessions, strong beta rhythms around 16 Hz could be observed as well as weaker rhythms around 32 Hz. In this study we focus on the beta rhythms and filtered the data between 12 and 20 Hz using a zero-phase 4-th order Butterworth bandpass filter.

**Figure 1:**
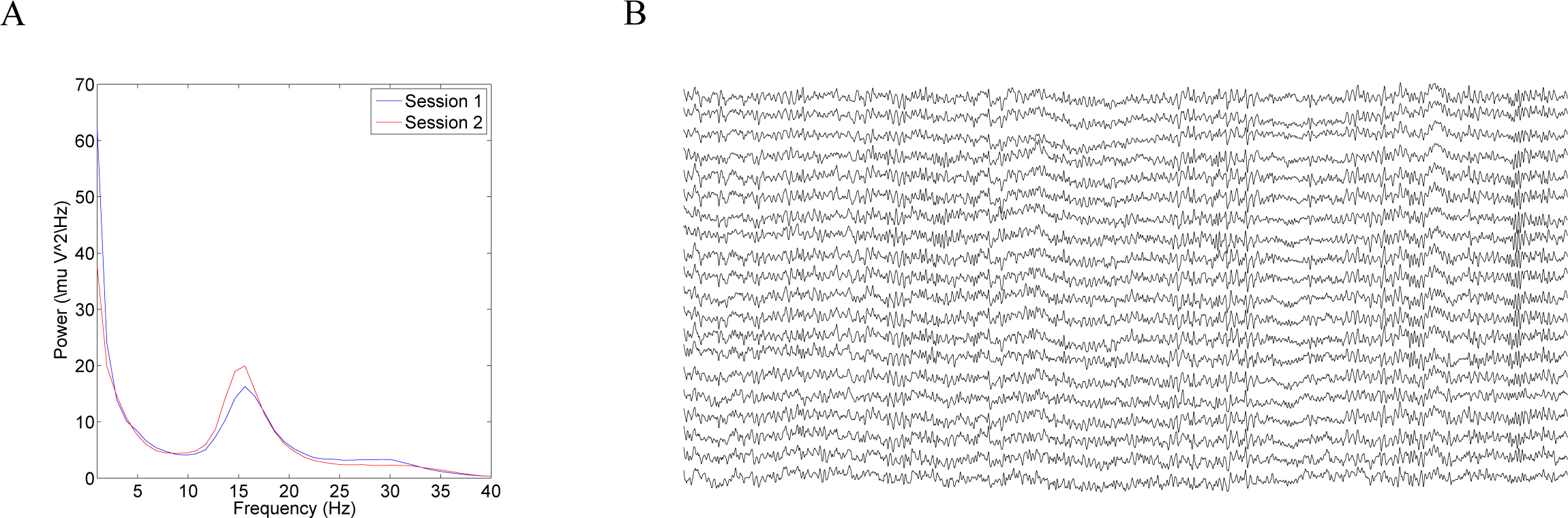
Electroencephalographic recordings. A. Electrode-averaged power spectra for each of the two five-minute recordings sessions (session 1 (blue) and session 2 (red)). B. Ten-second traces of the 20 most anterior electrodes. Note the strong presence of strong (≈ 16 Hz) beta oscillations and weaker (≈ 32 Hz) gamma rhythms.

### 2.2 Construction of ECoG leadfields

In order to construct a monkey-specific realistic ECoG leadfield matrix, the recording electrodes were localized relative to the monkey’s brain by the following procedure. During implantation surgery, pictures of the monkey brain were taken with a Nikon D5000 camera (42882848-pixel resolution, RGB color space) to document the position of the metal contacts relative to anatomical landmarks (sulci, gyri, vasa). The pictures, together with a triangulation of the cortical surface, were used as inputs to a custom-made interactive MatLab program to mark the electrode contacts. The triangulation of the cortical surface was obtained by segmenting a pre-surgical T1-weighted MRI (magnetization prepared rapid-gradient echo MPRAGE volumes acquired with Siemens tfl3d1_ns_ pulse sequence with flip angle = 8°; FOV = 18 cm, voxel size = 0.7 × 0.7 × 0.7 mm^3^, number of slices = 192, head first supine) into cerebrospinal fluid, white and grey matter using monkey-specific tissue priors [39] in SPM8. Lastly, the manually obtained electrode coordinates were corrected to achieve nominal inter-electrodes distance (2,5 or 3 mm according to the grid finger) and to follow geodesic curvilinear trajectories.

The source-space for the leadfield matrix was obtained from the T1-weighted MRI image using Caret software [55]. This yielded a high-resolution triangulation of the monkey’s cortical surface comprising 21339 vertices, which were used to define the source-space. Subsequently, a two-layer boundary-element method (BEM) realistic volume-conductor model (dura, white/gray matter boundary) was constructed using OpenMEEG [18] and FieldTrip [44]. The BEM yields the electric potential on the vertices of the outermost surface that are generated by unit-strength current dipoles located at the vertices of the cortical mesh, which is enclosed by both layers [43]. OpenMEEG requires four inputs: triangulated boundary surfaces of the head conductors with spherical topology, the position of the dipolar sources, the electrodes positions, and the conductivity values within the two compartments.

The outer boundary surface (i.e. the inner skull) was constructed by applying morphological operations (opening, closing, smoothing) and thresholding the segmented T1-weighted MRI volume, followed by triangulation [13]. Spherical topology was obtained by correcting the triangulation using custom geometry functions in FieldTrip. For numerical reasons related to calculation speed and intrinsic software limitations, we distributed no more than 10000 vertices on this surface. Vertex locations were chosen adaptively, which a denser coverage over the electrode grid. The inner boundary surface (i.e. the epicortical surface) was now obtained by in inward shift of the outer boundary surface by 1 mm. The conductivity values of the outer and inner compartments were set to 1 and 0.33 S/m, respectively [35]. The obtained leadfield matrix was subsequently converted to yield average-montage electric potentials at the 122 selected recording electrodes.

### 2.3 Source-space projection

The data were projected to the source-space using Harmony [46], which is a smoothed version of the classical minimum-norm estimate (MNE) designed to reduce surface-bias in the source reconstructions. Let *m* and *n* denote the number of recording electrodes and source-space locations (i.e. cortical vertices), respectively, and let *t* denote the number of samples. The 3*n* × *t*-dimensional vector source-activity matrix *J* is obtained by applying the Harmony inverse operator 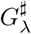 to the *m* × *t*-dimensional data matrix *X*:

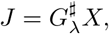

where

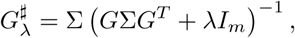

where *λ* ≥ 0 is the noise regularization parameter, the 3*n*-dimensional matrix Σ denotes the *a priori* covariance matrix of *J, I_m_* denotes the m-dimensional identity matrix, and T denotes matrix transpose. Harmony models Σ as

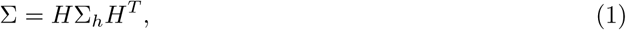

where *H* denotes the 3*n* × 6(*l_max_* + 1)^2^-dimensional Harmonic transformation matrix, which contains the spherical harmonics up to and including degree *l_max_* = 20. The 6(*l_max_* + 1)^2^-dimensional matrix Σ_*h*_ denotes the *a priori* source covariance matrix expressed in the Harmonic basis and is modeled as a diagonal matrix with diagonal entries equal to 1/(1 + *l^p^*), where *p* = 0.5 controls how fast the *a priori* source variance decreases with degree and therefore indirectly controls the level of spatial smoothing in the source reconstructions [46]. The spherical harmonics are calculated on the vertices of a triangulation of the spherical cortical surface that is induced by the triangulation of the fiducial cortical surface using Caret software [55]. After source-space projection, the vector-valued time-series at each cortical vertex was reduced to a scalar-valued time-series by projecting it onto its first eigenvector.

Appropriate values for *λ* were determined by applying the *L*-tangent norm criterion [4] to 100 randomly chosen samples and averaging the obtained values. The *L*-tangent norm criterion selects the value of *λ* that minimizes the speed along the (logarithm of the) *L*-curve. Let

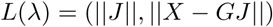

be the *L*-curve [23], then *λ* is chosen as follows:

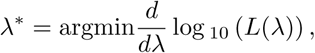

which was obtained by calculating the *L*-curve on the (logarithm of the) interval [−6, 0] in steps of 0.1 and by approximating *d/dλ* in these points by (forward) finite differences.

### 2.4 Reconstruction of functional networks

After obtaining the source-space scalar time-series, the generators (i.e. network nodes) are identified using spatial independent component analysis (ICA). Spatial ICA is appropriate in this context, because the number of samples *t* is an order of magnitude less than the number of cortical vertices *n*. The appropriate number of components *k* is determined by inspection of the eigenspectrum of the sensor-level data matrix. Before applying ICA, the rank of data matrix was reduced by retaining only the first *k* eigenvectors. This was done because in our simulations, this improved the identification of generators (results not shown). Because in practice, it is not always clear how to set *k*, we performed simulations in which *k* was set to a different value than the true number of generators. Spatial ICA is used only to estimate the number of generators and to reconstruct their spatial profiles because their associated time-courses generally not well resemble the true generator time-series (results not shown). Reconstructed generator time-courses are obtained by selecting the source-level time-course of the cortical voxel that correlates most strongly with the generators’ ICA time-course.

Studies on source-level resting-state networks using MEG (and EEG) typically focus either amplitude envelope correlations [42] or on oscillatory phase-locking or coherence [26]. We therefore consider the reconstruction of both types of functional connectivity: amplitude correlation and oscillatory phase-locking. Let *s*_1_(*t*) and *s*_2_(*t*) be the reconstructed time-courses of two identified generators. The *amplitude correlation ρ* between *s*_1_(*t*) and *s*_2_(*t*) is defined as the Pearson correlation between the instantaneous amplitudes *a*_1_(*t*) and *a*_2_(*t*) of *s*_1_(*t*) and *s*_2_(*t*), which are defined as

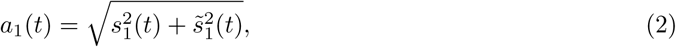

where 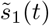 is the Hilbert transformation of *s*_1_(*t*) and similarly for *s*_2_(*t*). The amplitude correlation ranges between −1 and 1 and measures the strength of the instantaneous linear dependence between *a*_1_ and *a*_2_. Oscillatory phase-locking between *s*_1_ and *s*_2_ is measured by the complex-valued *phaselocking-value ν*, which is defined as

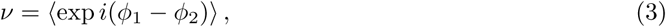

where *ϕ*_1_(*t*) and *ϕ*_2_(*t*) denote the instantaneous phases of *s*_1_(*t*) and *s*_2_(*t*), and the brackets denote temporal averaging. The phase-locking value *ν* is complex-valued, where its absolute value, which ranges between 0 and 1, measures the strength of oscillatory phase-locking between *s*_1_ and *s*_2_ and its phase measures their phase-lag (i.e. time-delay).

### 2.5 Simulated data

The proposed reconstruction method was tested by simulating ECoG data comprising a network of *k* spatially extended cortical generators. The center location of the *i*-th generator is denoted by *x_i_* and its amplitude at location *x* as modeled as

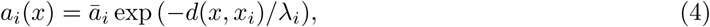

where 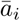 denotes the generators’ (maximal) amplitude, *λ_i_* its characteristic scale, and *d*(*x*, *x_i_*) the geodesic distance between *x* and *x_i_* along the cortical surface. The time-course of the *i*-th generator was simulated by generating a Gaussian white noise signal, bandpass filtering it ±2 Hz around the generators’ peak frequency *f_i_*, and subsequently normalizing it to variance one. Because the time-course is generated independently for each generator, this yields non-phase-locked oscillatory time-series a* with uncorrelated amplitude envelopes. Correlations are incorporated by mixing the generators’ time-courses according to

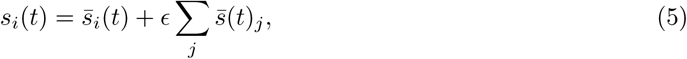

where ϵ ≥ 0 is the mixing coefficient that controls the overall correlation, and where *j* runs over all generators except the *i*-th. The mixed time-series were subsequently normalized to variance one. This yields a network of oscillatory generators which are phase-locked at zero lag and that whose amplitudes are instantaneously correlated. The level of phase-locking and amplitude correlation are determined by the value of *ε*. Lagged phase-locking was incorporated by translating the generators’ time-series over *τ_k_* milliseconds, relative to the first generator. Simulation of this network returns yields *n* × *t*-dimensional matrix *S*, with *n* the number of cortical vertices and *t* the number of samples, containing the simulated time-series in its rows.

The simulated data were subsequently projected to the ECoG grid and measurement noise was added, yielding a simulated data matrix *X*:

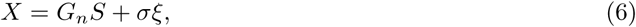

where *G_n_* denotes the scalar leadfield matrix, which is derived from the vector leadfield matrix by assuming the sources to be perpendicular to the cortical surface, and *ξ* denotes Gaussian noise with covariance matrix taken from the empirical data, filtered between 60 and 80 Hz, and scaled to have maximal value one. Since this frequency band still contains physiological signal, *ξ* models a combination of spontaneous cortical background activity and sensor noise. The noise intensity *σ* was chosen to obtain a signal-to-noise ratio (SNR) of *η* (a free parameter), and in its calculation the signal power was taken to be the average power across electrodes.

We placed *k* = 5 generators at locations (partially) underneath the electrode grid and will refer to these as generators 1, 2, 3, 4, and 5. Their spatial profiles are shown in Figure 2A. Generator 1 is located on the posterior wall of the central sulcus, generators 2 and 5 in the supplementary motor area, just outside the grid, generator 3 on the wall of the superior arcuate sulcus, and generator 4 on the crown of the anterior central gyrus. To make the network realistic and challenge the reconstruction method, the generators’ amplitudes, characteristic scales, peak-frequencies, and time-lags (relative to the first source) differed between generators and are listed in Table 1. Observation time and sampling frequency were equal to those of the experimental data (5 minutes and 500 Hz, respectively). For each choice of the parameters, we averaged the results over ten realizations. Figure 2B shows ten-second epochs of the simulated ECoG data.

**Figure 2:**
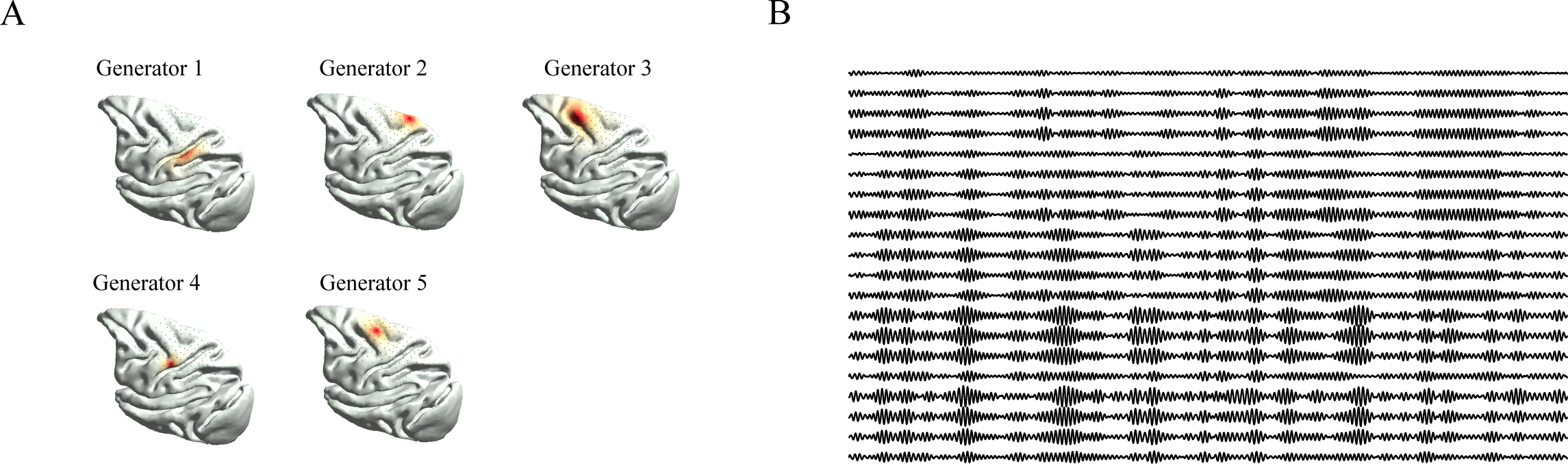
Simulated ECoG data. A. Amplitude distribution of the five simulated cortical generators scattered throughout the somatosensory, motor, and premotor cortices. The black dots denote the locations of the registered ECoG electrodes. B. Ten-second epoch of simulated ECoG data. The signals of the 20 most anterior electrodes are shown.

**Table 1:** Simulation parameters. Listed are the parameters, their symbols, and values/ranges. The subscript *i* refers to the *i*-th generator. If the third column contains a single value for a given parameter, the value applies to all *k* generators.

| Parameter | Symbol | Values/range |
| --- | --- | --- |
| Number of generators | $k$ | 5 |
| Maximal amplitude | $\bar{a}_i$ | 2, 3, 1, 4, 2 |
| Characteristic scale (mm) | $\lambda_i$ | 4, 3, 5, 2, 3 |
| Peak-frequency (Hz) | $f_i$ | 16, 15, 17, 18, 16.5 |
| Mixing coefficient | $\epsilon$ | 0.5 |
| Time-lag (ms) | $\tau_i$ | 10, 20, 30, 0 |
| Signal-to-noise ratio (dB) | $\eta$ | 5, 10, 15, 20, 25 |

## 3 Results

### 3.1 Identification of cortical generators

Before we discuss the simulation results, we motivate the use of spatiotemporal decomposition methods like ICA, by showing that it is generally difficult to uncover the true network structure by analyzing either the spatial or the temporal structure of the reconstructed activity. The former approach is referred to as *source-amplitude imaging* [30] and the latter to *seed-based functional connectivity*. The latter approach is frequently used to reconstruct functional networks from source-projected MEG (and EEG) data. Figure 3A shows the distribution of reconstructed cortical amplitudes from a single realization of 10 dB simulated data. Observe that although two generators are clearly visible (generators 2 and 4), it is difficult to infer exactly how many more generators there are: it can range anywhere between two and four. In particular, the existence of generator 3 would not even be suspected. Thus, by itself, and although useful, source-amplitude imaging is generally insufficient to identify the true number of generators. Figure 3B shows the covariance map of the band-limited amplitude fluctuations with seed chosen at the location of (reconstructed) generator 2. Although its connectivity to generator 4 is visible, its connectivity to the other generators is missed. This example motivates an approach that integrates the spatial and temporal structure in the source-projected data.

**Figure 3:**
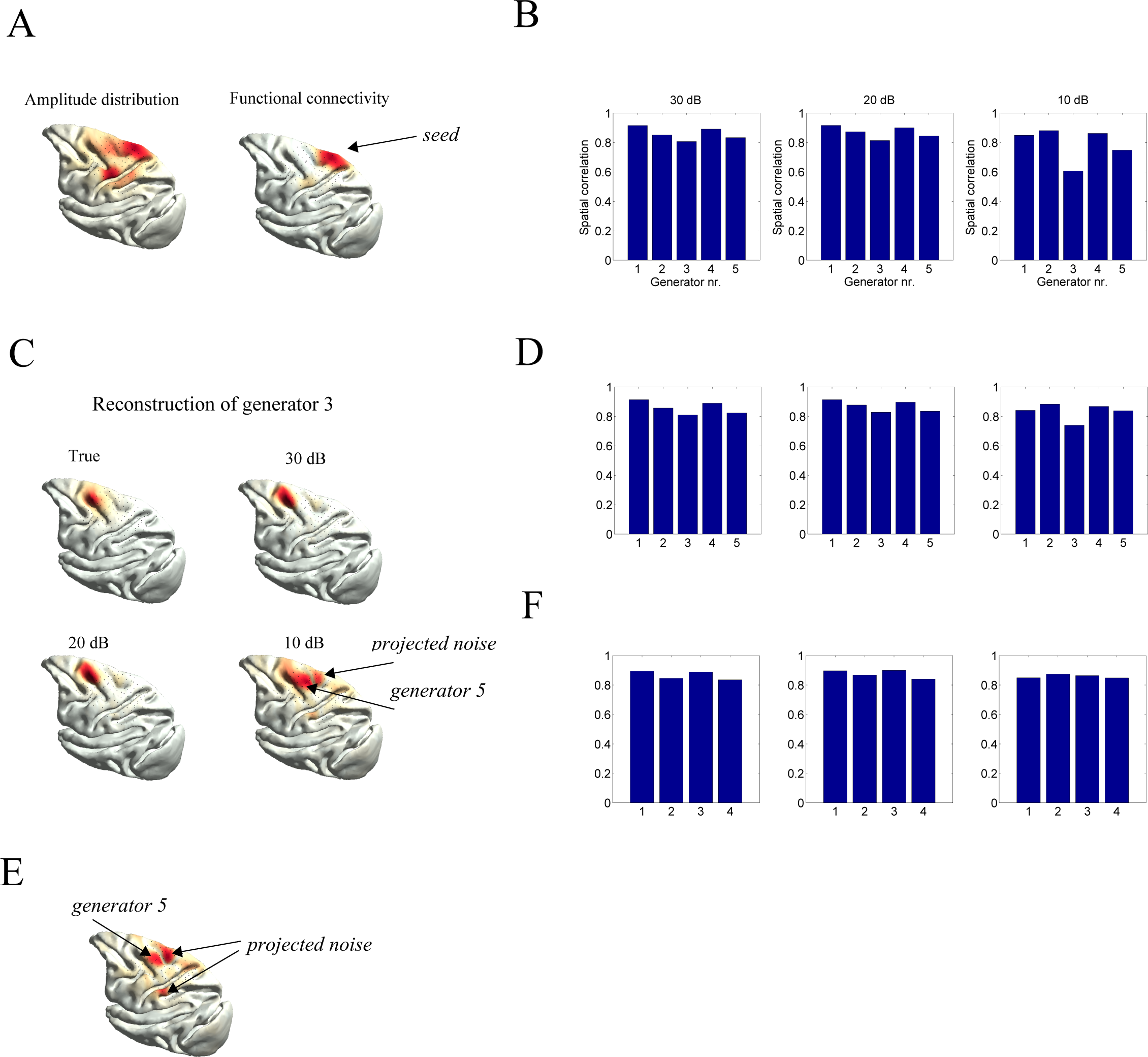
Delineation of cortical generators. A. Reconstructed distribution of cortical amplitude for a single realization of simulated data with a SNR of 10 dB (left) and seed-based covariance map with seed at the location of generator 5 (right). B. Spatial correlation coefficients between the true and reconstructed generators for all three noise-levels (10, 20, and 30 dB) for the case that the number of independent components was set to its true value (*k* = 5). C. True amplitude map of generator 3 and its reconstruction for all three noise-levels (10, 20, and 30 dB). D. Spatial correlation coefficients between the true and reconstructed generators for all three noise-levels (10, 20, and 30 dB) for the case that the number of independent components was set too high (*k* = 6). E. Spatial profile of the sixth independent component. It comprises a mixture of (true) generators and projected sensor-noise. The spatial profiles and correlation coefficients were obtained by averaging over ten realizations. F. Spatial correlation coefficients between the true and reconstructed generators for all three noise-levels (10, 20, and 30 dB) for the case that the number of independent components was set too low (*k* = 4).

The eigenspectra of the simulated ECoG data show that *k* = 5 is an appropriate choice for the number of generators (i.e. independent components). For comparison with the true generators, the reconstructed generators were ordered to match the true generators, based on their spatial correlations. For all three noise-levels, the true generators could be unambiguously paired with the true generators. Figure 3B shows the spatial correlation coefficients between the true and reconstructed generators for the three noise-levels. The correlations are generally high and roughly equal for all noise-levels, except for generator 3, which is less well reconstructed in case of noisy data (10 dB). This is due to the relative low amplitude of this generator (Table 1). Figure 3C shows the amplitude profile of generator 3 and its reconstructions for al three noise-levels. Although generator 3 is well reconstructed for low and mid levels of noise (20 and 30 dB), for noisy data (10 dB) it becomes mixed with other generators (in particular with nearby generator 5) and with projected sensor-noise (designated by the arrow in Figure 3C). These simulations demonstrate that ICA is able to identify cortical generators based on their band-limited amplitude fluctuations, but that weak generators might be inaccurately reconstructed for noisy data. Unfortunately, their is *a priori* no way of telling whether this is the case, since this depends on the combination of several factors such as the spatiotemporal characteristics of the sensor-noise, the forward model, and the organization of cortical activity, which is unknown.

In practice, it is not always clear what the appropriate number of independent components is and it is therefore relevant to consider the effect of setting *k* to a value higher or lower than the true number of generators. We therefore repeated the simulations while setting *k* = 6 and *k* = 4. Figure 3D shows the spatial correlations between the five true generators and the five unambiguously paired reconstructed generators. The correlations are roughly equal to those obtained above by setting *k* = 5, which means that setting *k* too high does not distort the reconstruction of the true generators. Perhaps surprisingly, the spatial map of generator 3 is more accurately reconstructed than when setting *k* to its true value (*k* = 5). Figure 3E shows the spatial map of the sixth independent component clarifies why this in the case: instead of mixing generator 5 and projected sensor noise into reconstructed component 3, they together form the sixth independent component. When the number of generators is set too low (*k* = 4), the weakest generator (generator 3) is missed, but the other generators are correctly identified and their spatial correlations with the true generators remain high (Figure 3F).

There thus appears to be an advantage in setting *k* too high as this tends to unmix the true generators by grouping generator copies and projected sensor-noise into a new generator. This advantage can often be gained when using other decomposition methods such as (*k*-means) clustering algorithms and (non-negative) matrix factorization. The disadvantage, however, is that in practice, the true number of generators is generally unknown, and it might therefore not be possible to decide which generator is spurious. An ad hoc criterion that happens to work in the simulations above is to visually inspect the spatial profiles of the reconstructed generators. The spurious generator is then set apart from the others by its composite nature (Figure 3E) and overlap with other generators. There is, however, no guarantee that this criterion is generally valid. From a practical perspective, setting the number of generators too low is less disadvantageous because although generators might be missed, the spatial profiles of the identified generators remain accurate. This then allows for the accurate reconstruction of a subnetwork of the true functional network.

### 3.2 Reconstruction of band-limited amplitude correlations

Figure 4A shows the true and reconstructed amplitude correlation matrices for the three noise-levels. For mid and low noise-levels (20 and 30 dB) the amplitude correlation matrices are accurately reconstructed, while for noisy data (10 dB), the functional connectivity map of generator 3 is not well reconstructed. Specifically, its connectivity to generator 5 and generator 1 are over- and underestimated, respectively. Because the generators’ time-courses are extracted based on their reconstructed spatial profiles, (in)accurate reconstruction of functional connectivity depends entirely on the accuracy of these reconstructions. In the previous section we have seen that for noisy data (10 dB), reconstructed generator 3 is mixed with generator 5 (as well as with projected sensor-noise), which causes the overestimation of their connectivity. Mixing of generator 5 into reconstructed generator 3 is also responsible for the underestimation of its connectivity with generator 1, as the connectivity with generator 1 and 5 is weaker than that with generator 3.

**Figure 4:**
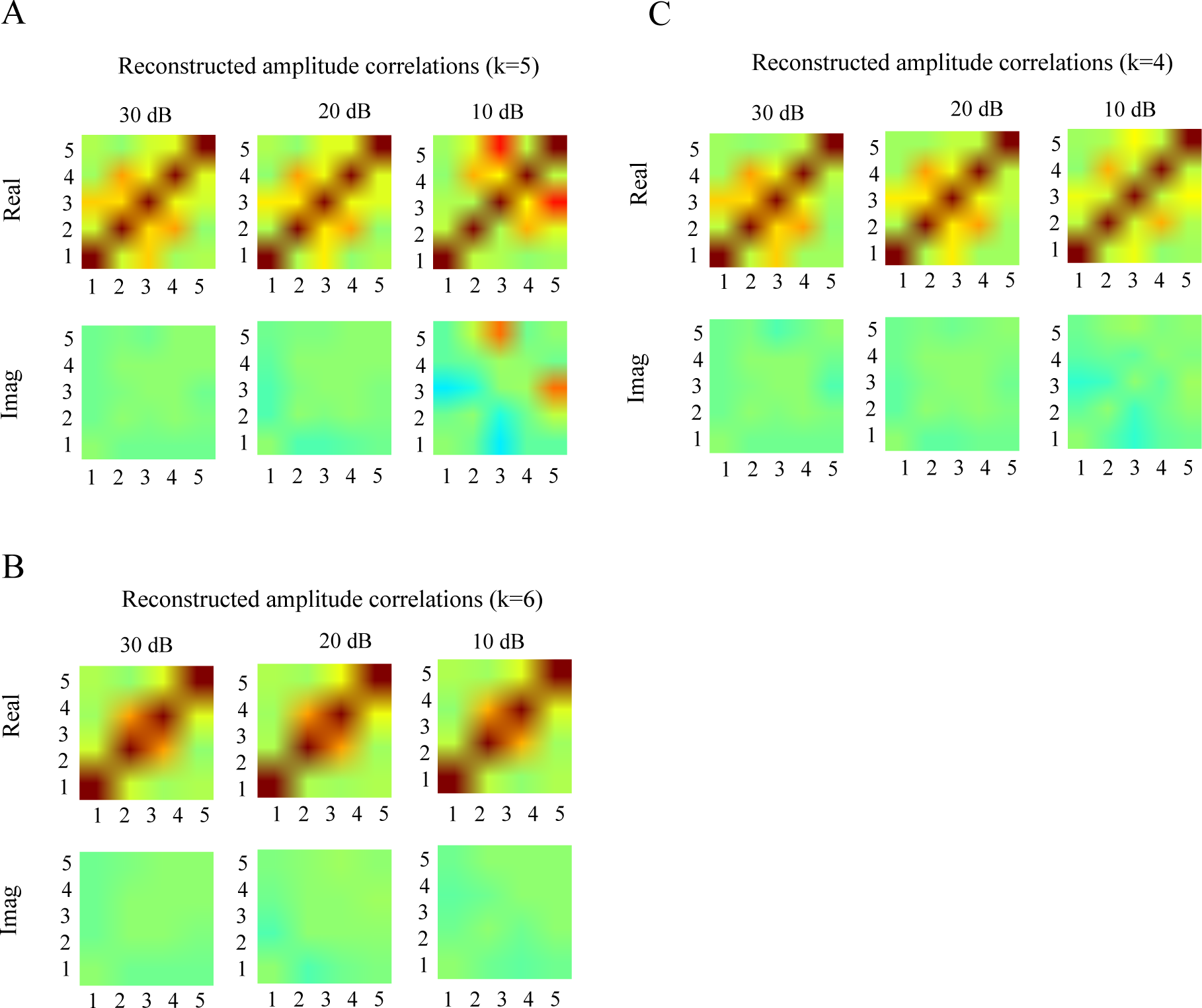
Reconstruction of amplitude envelope correlations. A. Top row: true and reconstructed amplitude envelope correlations matrices for all three noise-levels (10, 20, and 30 dB). Bottom row: Differences between the true and reconstructed amplitude envelope correlation matrices for all three noise-levels. The number of independent components was set to *k* = 5. B and C: Same format as A. but with the number of independent components set to *k* = 6 and *k* = 4, respectively.

In the previous section, we have seen when the number of independent components is set too high (*k* = 6), all generators can still be identified and that for noisy data (10 dB), the reconstructed spatial profiles are in fact more accurate, as compared to setting *k* to the true number of generators (*k* = 5). As a consequence, the reconstructed functional connectivity matrices also become more accurate, because the amplitude envelopes used for their reconstruction are based on the spatial profiles of the reconstructed generators. This is shown in Figure 4B. In particular, for noisy data, the connectivity of generator 3 with generators 1 and 5 is now accurately reconstructed. In practice, however, this is of limited use because the true number of generators is unknown and there is no general way of deciding which reconstructed generator is spurious and should be discarded from the analysis. In the previous section we have also seen that setting the number of components too low (*k* = 4), yields accurate spatial profiles of four of the five generators, although the weakest generator (generator 3) is missed. Figure 5 shows that this yields accurate reconstruction of the subnetwork comprised of generators 1, 2, 4, and 5.

### 3.3 Reconstruction of oscillatory phase-locking

Figure 5A shows the true (black lines) and reconstructed (complex-valued) phase-locking factors between all generator pairs and for all three noise-levels (blue, red, and green lines correspond to 30 dB, 20 dB, and 10 dB, respectively) for the case *k* = 5. The phase-locking factors have been restricted to the first quadrant by taking the absolute values of its real and imaginary parts. This is done because the signs of the basis vectors in which the leadfields are expressed, are arbitrary and therefore do not allow the signs of the source time-courses to be determined. This means that the phase-locking factors can be determined only modulo 180 degrees, unless the local orientations of the cortical surface are used. Note that the phase-locking factors are well reconstructed for 20 and 30 dB data, while for 10 dB data, the reconstructions are less accurate, at least for some pairs of generators. We make several remarks about the 10 dB case. First, the phase-locking magnitudes can be under-as well as overestimated. Second, the relative phase-differences, that is, the angles of the phase-locking factors with the x-axis, are underestimated for most pairs of generators. This effect is referred to as *phase-contraction* and refers to a reduction in relative phases due to volume conduction [27]. Third, and in contrast to amplitude correlations, inaccuracies in the reconstruction of phase-locking factors are not limited to generator 3 and 5, but affect most pairs of generators. This reflects the fact that phases are more susceptible to volume conduction than amplitudes [27]. Fourth, although the relative phase-differences are underestimated for most pairs of generators, the some pairs, the phase-differences are overestimated. This is the case for generator pairs (1,3), (1,4), and (4,5). In particular, while these generator pairs of coupled instantaneously i.e. with phase-difference zero, their reconstructed imaginary pairs are non-zero. Although imaginary (i.e. lagged) phase-locking or coherence can never arise from a pair of non-interacting generators - which has motivated the use of lagged interaction measures [40, 53], this is only true in the absence of lagged generators [10]. This second-order effect of volume conduction can be witnessed in our simulations and warrants care when applying lagged interaction measures. When the number of independent components is set to a different value than the true number of generators (*k* = 6 or *k* = 4), the accuracy of the reconstructed phase-locking matrices improves, as is the case with amplitude correlation matrices (see Section 3.2). The reconstructed phase-locking matrices still suffer from phase-contraction, which again reflects the relative susceptibility of phases for volume-conduction.

**Figure 5:**
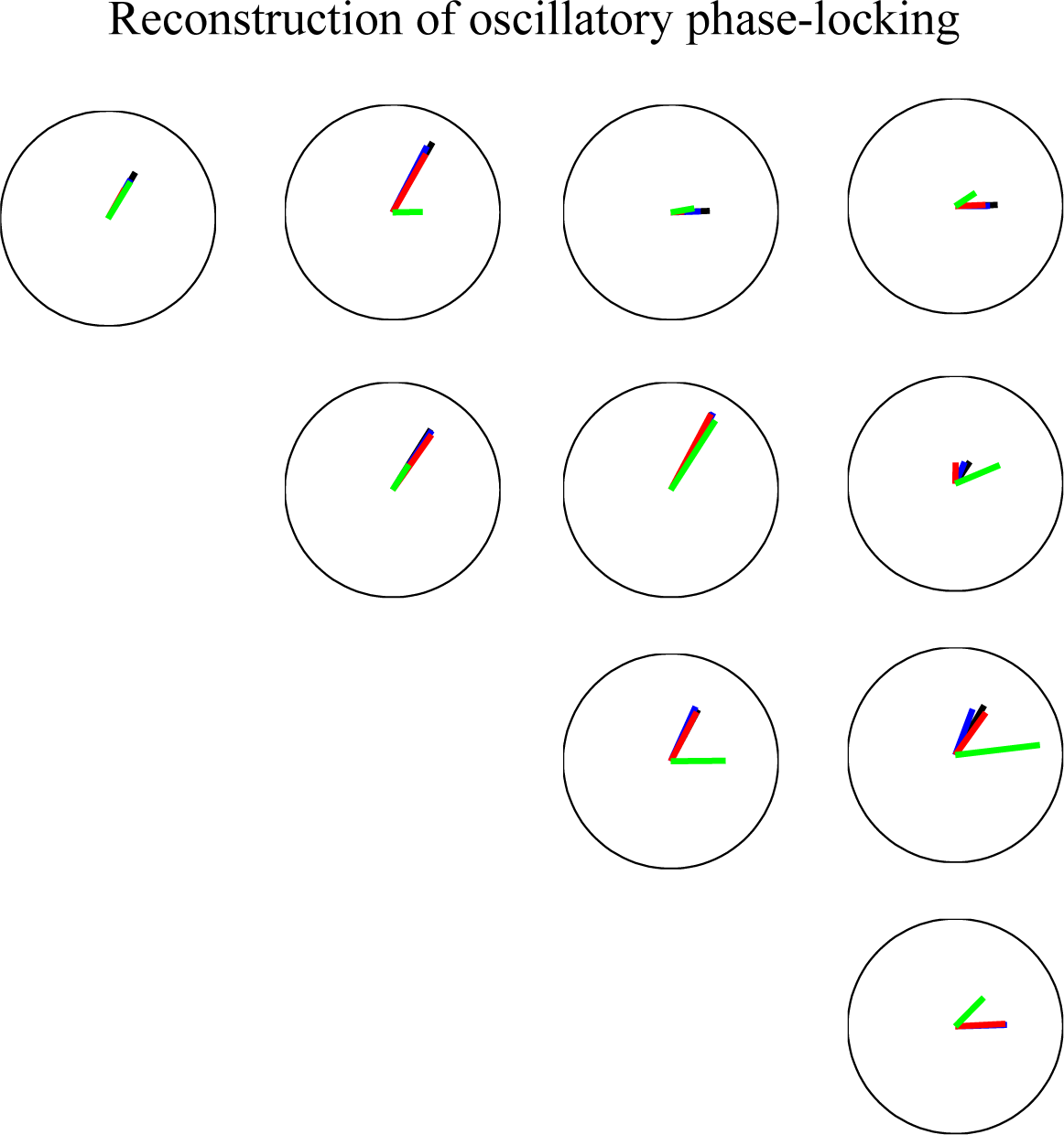
Reconstruction of oscillatory phase-locking. True (black) and reconstructed complex-valued phase-locking factors between all pairs of generators and for three signal-to-noise levels. Blue, red, and green correspond to 30 dB, 20 dB, and 10 dB, respectively. The number of independent components was set to *k* = 5.

### 3.4 Sensorimotor beta oscillations in macaque neocortex

In this section we apply the reconstruction method to macaque beta oscillations in the sensorimotor system recorded with ECoG (see Section 2.1). Figure 6A shows the eigenspectra of the sensor-data covariance matrices of both recording sessions. A reasonable choice for the number of independent components appears to be *k* = 7. Figure 6B shows the spatial profiles of the reconstructed generators. Two of the generators are located in the primary motor cortex on the pre-central gyrus (generators 1 and 2), two in the pre-motor cortex (generators 3 and 4), and three in the somatosensory cortex on the post-central gyrus and in the intra-parietal sulcus (generators 5, 6, and 7). Note that the generators are spatially extended, which confirms recent multi-electrode recordings of beta oscillations in macaque primary and pre-motor cortices [50, 54]. Although their spatial extent might be overestimated to some degree, which is due to the noise-regularization term in the inverse solution, these observations are surely incompatible with point sources i.e. equivalent current dipoles. Also note that, taken together, the reconstructed generators cover the post-central gyrus over its entire lateral extent and most of the pre-motor cortex. These data therefore suggest that sensorimotor beta oscillations are a spatially continuous phenomenon, rather than a finite network of discrete generators.

**Figure 6:**
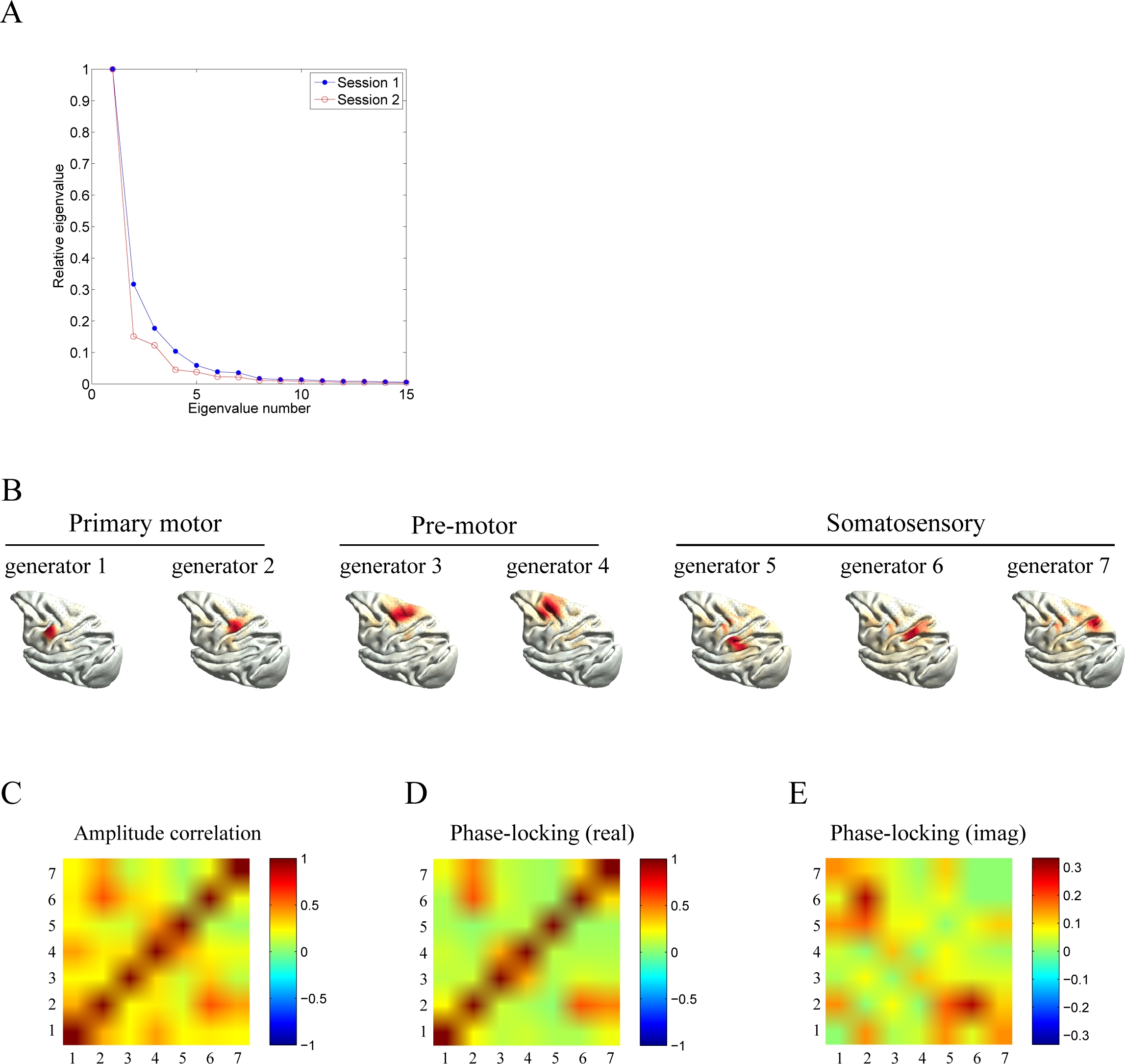
Functional organization of sensorimotor beta oscillations in macaque cortex. A. Eigenvalues of the sensor-data covariance matrices of session 1 (blue) and session 2 (red). B. Reconstructed cortical beta generators. C. Correlation matrix of band-limited amplitude fluctuations between the reconstructed generators. D. Real part of the phase-locking matrix between the oscillatory time-courses reconstructed generators. E. imaginary part of the phase-locking matrix between the oscillatory time-courses reconstructed generators. Note that the color-scaling in panel E is different than that used in panels C and D.

Figure 6C shows the correlation matrix between the band-limited amplitudes of the reconstructed generators and Figures 7D and E show their (absolute) real and imaginary phase-locking matrices. All correlations are significant (*p* < 0.01, Bonferroni-corrected) which was tested by approximating the corresponding null-distributions using phase-randomized surrogate data [45]. We make three observations. First, the strongest correlations are between generator 2, 5, and 6. One possible explanation for this is that the number of independent components is set too high and these three generators actually comprise a single generator that has been broken up into three parts due to projected sensor-noise. Indeed, when the number of independent components is set to *k* = 5, generators 5 and 6 get merged (and generator 4 disappears). As we will see below, these three generators also have high imaginary phase-locking factors, which means that their oscillations are lagged with respect to each other. We therefore argue that they reflect a *traveling wave* that has been broken up into locally coherent generators. Another explanation for the high amplitude correlation between generator 2 and 6 is the existence of a single generator at the fundus of the central sulcus that has (erroneously) been split into two components. If this were the case, however, the oscillations of these generators would be locked with zero phase, while in fact they have a consistent and large phase-difference. We take this as evidence for the existence of a traveling wave that propagates through the central sulcus. Second, there are generators that are spatially well separated but nevertheless have relatively strong amplitude correlations. This is the case for generator-pairs (1,4), (2,7), and (4,5). Third, most generator pairs have non-zero imaginary phase-locking matrices, which means that their oscillations are locked with non-zero lag. Although the exact values might be distorted by (first and higher-order) volume conduction effects, such a scenario, together with the spatial extent of the generators, points towards complex spatiotemporal dynamics including wave propagation.

## 4 Discussion

In this study, we have assessed the feasibility of delineating cortical generators from oscillatory ECoG data and reconstructing their functional interactions. In particular, cortical generators were delineated by applying spatial independent component analysis (ICA) to source-projected ECoG data. We considered two kinds of functional interactions: (instantaneous) correlations between band-limited amplitude envelopes and oscillatory phase-locking. While amplitude correlations are widely used in EEG and MEG resting-state studies [9, 52, 34] and follow the correlation patterns obtained from functional magnetic resonance imaging (fMRI) [38, 29], oscillatory phase-locking yields dynamical information on a much finer time-scale and therefore makes full use of the high temporal resolution of electrophysiological recordings. Using a simulated network of five cortical generators, we showed that amplitude correlations and oscillatory phase-locking can both be accurately reconstructed, although the latter generally requires a higher signal-to-noise ratio. We applied the methodology to resting-state recordings of a macaque monkey displaying strong sensorimotor beta (≈ 16 Hz) oscillations and found these to constitute a set of coordinated generators covering primary and second-order somatosensory cortices, primary motor cortex, and pre-motor regions. Our study thus shows that ECoG recordings allow for a more detailed characterization of resting-state cortical oscillations than what can be obtained with non-invasive recordings (EEG/MEG) and opens up the possibility of studying oscillatory cortical dynamics on a spatial scale that lies between that of EEG/MEG and intra-cortical LFP recordings.

Our simulations suggest that band-limited power-fluctuations of cortical generators and their correlations can be reasonably well reconstructed from ECoG channel data, even for noisy data (10 dB). Reconstruction of oscillatory phase-locking is accurate as well, as long as the data are not too noisy. It might be suspected that the problem lies in the estimation of the (unknown) dipole orientations. This is not the case, however, as the reconstructed phase-locking matrices remain practically unchanged when the dipole orientations are assumed to be known (results not shown). Rather, phases are more susceptible to volume conduction than amplitudes [27] and, as such, are more difficult to reconstruct. We have not explicitly evaluated the performance of lagged phase-locking measures such as the phase-lag index [53], which are designed to measure interaction in the presence of volume conduction. Instead, we have analyzed how volume conduction distorts the reconstructed (complex-valued) phase-locking factors as this provides a reference for interpreting the results obtained from using such measures. For instance, we found that due to second-order effects of volume conduction [10], reconstructed lags between pairs of generators can have both a positive and a negative bias, and can lead to non-negative lags while in reality, the generators are coupled with zero lag. This unfortunate situation makes it difficult to interpret observed values of lagged interaction measures in terms of interaction strengths and it seems unavoidable to develop measures that explicitly deal with higher-order effects [10].

To project the channel data to the cortical surface, we have used a smoothed version of the minimum norm estimate (MNE) [46]. We have also experimented with normalized inverse methods such as dynamic statistical parametric mapping (dSPM), standardized low resolution electromagnetic tomography (sLORETA), and weighted minimum norm estimate (wMNE) [19], which all yielded less accurate source reconstructions. The reason from this is that ECoG leadfields have a much broader range (as measured by their norms) than EEG and MEG leadfields, due to the limited coverage of the electrode grid. As a consequence, the source reconstructions are typically biased towards cortical locations that are far from the grid. These findings are in line with another ECoG simulation study, which found MNE to outperform both sLORETA and WMNE [6]. In another study, however, WMNE was found to yield reasonable reconstructions [56], although no comparison with other methods was carried out. In [12], the Multiple Signal Classification (MUSIC) algorithm was found to perform well, although it was not directly compared with other methods. Another option are adaptive spatial filters (beamformers) [19]. Unlike in the case of EEG and MEG data, in which the dipole approximation might be reasonable, the close proximity of ECoG electrodes to the cortex requires simulation of spatially extended sources (as in our study), at least in case of resting-state data, which is dominated by strong oscillatory activity. The resulting local correlations might prevent successful application of beamformers to ECoG data. As it is not the goal of this study to carry out a exhaustive comparison of different inverse methods, a comparative analysis, and adaptation, of EEG/MEG inverse methods to ECoG data is certainly an important research direction.

The reconstruction method provides evidence that resting-state beta oscillations are widespread throughout the somatomotor system. In particular, they cover post-central gyrus and the walls of the postcentral sulcus over their entire lateral extents (primary somatosensory cortex), the intra-parietal sulcus (higher-order somatosensory cortex), the pre-central gyrus and the walls of the pre-central sulcus (primary motor cortex), and extend into more anterior regions, including the lateral end of the arcuate sulcus (premotor cortex). The generation of beta oscillations within the primary motor cortex (precentral gyrus) is well-established and can also be observed non-invasively in humans with EEG and MEG [24, 31, 48, 5]. The existence of beta generators in frontal (pre-motor) regions is also well-established from invasive (LFP) recordings in macaque monkeys [51], but is not commonly reported in human EEG and MEG studies and the same is true for beta oscillations in post-central regions [3]. ECoG source modeling hence yields a more detailed impression of the distribution and number of cortical beta generators within the sensorimotor system than EEG and MEG do and resemble more the findings reported in LFP studies. ECoG recordings are also advantageous over intra-cortical LFP recordings as they allow for a more macroscopic picture (order of centimeters) of the cortical organization of cortical oscillations.

Our study suggests that beta oscillations comprise a distributed functional network that covers both primary and higher-order motor and somatosensory cortices. This is in line with local field potential (LFP) recordings from multiple intra-cortical electrodes located in pre- and post-central regions [51, 3]. In particular, in [3], directed oscillatory coupling was reported between beta oscillations within the primary and pre-motor cortices and somatosensory cortex, including the intra-parietal sulcus. In our study we did observe relatively large latencies between the beta oscillations originating from pre- and post-central regions, which is at least consistent with the directional coupling from somatosensory to motor cortex observed in [3]. As noted above, (source-projected) ECoG data also has advantages over intra-cortical LFP recordings in that they do not suffer from sparse sampling from the cortical surface. As such, they allow a description of cortical oscillations that goes beyond the discrete and point-like characterization offered by intra-cortical LFP recordings. In particular, when the spatial profiles of the delineated generators (see Figure 7B) are combined, the oscillations appear to cover almost the entire sensorimotor system. This raises the question if the functional organization of cortical beta is best conceptualized as a network of discrete generators or as a spatially continuous system. In this context, it is interesting to note that intra-cortical microelectrode recordings in macaque primary motor cortex [54] and dorsal premotor cortex [50] have shown that at the scale of several millimeters, beta oscillations behave as traveling waves. Source-projected ECoG recordings possibly allow to scale up these and similar observations to the centimeter scale and provide a wider window into the spatiotemporal dynamics of cortical beta oscillations. This, however, requires accurate estimation of instantaneous oscillatory phases with high spatial resolution, and it is unknown to which extent existing inverse methods enable this.

## References

[1] Y. Aoki, R. Ishii, R.D. Pascual-Marqui, L. Canuet, S. Ikeda, M. Hata, K. Imajo, H. Matsuzaki, T. Musha, T. Asada, M. Iwase, and M. Takeda. Detection of EEG-resting state independent networks by eLORETA-ICA method. Front. Hum. Neurosci., 9(February):31, 2015.

[2] M.J. Brookes, M. Woolrich, H. Luckhoo, D. Price, J.R. Hale, M.C. Stephenson, G.R. Barnes, S.M. Smith, and P.G.q Morris. Investigating the electrophysiological basis of resting state networks using magnetoencephalography. Proc. Natl. Acad. Sci. U. S. A., 108(40):16783–8, oct 2011.

[3] A. Brovelli, M. Ding, A. Ledberg, Y. Chen, R. Nakamura, and S.L. Bressler. Beta oscillations in a large-scale sensorimotor cortical network: directional influences revealed by Granger causality. Proc. Natl. Acad. Sci. U. S. A., 101(26):9849–54, 2004.

[4] F. Brunet, A. Bartoli, and L. Uda. L-tangent norm: A low computational cost criterion for choosing regularization weights and its use for range surface reconstruction. Proc. Fourth Int. Symp. 3D Date Process. Vis. Transm., 2008.

[5] D.O. Cheyne. MEG studies of sensorimotor rhythms: A review. Exp. Neurol., 245:27–39, 2013.

[6] J. H. Cho, S. B. Hong, Y. J. Jung, H. C. Kang, H. D. Kim, M. Suh, K. Y. Jung, C.H. Im, and Chang-Hwan Im. Evaluation of algorithms for intracranial EEG (iEEG) source imaging of extended sources: feasibility of using iEEG source imaging for localizing epileptogenic zones in secondary generalized epilepsy. Brain Topogr., 24(2):91–104, jun 2011.

[7] J. H. Cho, H. C. Kang, Y.-J. Jung, J.-Y. Kim, H. D. Kim, D. S. Yoon, Y. H. Lee, and C. H. Im. Localization of epileptogenic zones in Lennox-Gastaut syndrome using frequency domain source imaging of intracranial electroencephalography: a preliminary investigation. Physiol. Meas., 34(2):247–63, feb 2013.

[8] J. S. Damoiseaux, S. A. R. B. Rombouts, F. Barkhof, P. Scheltens, C. J. Stam, S. M. Smith, and C. F. Beckmann. Consistent resting-state networks across healthy subjects. Proc. Natl. Acad. Sci., 103(37):13848–13853, 2006.

[9] F. de Pasquale, S. Della Penna, A.Z. Snyder, C. Lewis, D. Mantini, L. Marzetti, Paolo Belardinelli, Luca Ciancetta, Vittorio Pizzella, Gian Luca Romani, and Maurizio Corbetta. Temporal dynamics of spontaneous MEG activity in brain networks. Proc. Natl. Acad. Sci. U. S. A., 107(13):6040–6045, 2010.

[10] M. Drakesmith, W. El-Deredy, and S. Welbourne. Reconstructing coherent networks from electroencephalography and magnetoencephalography with reduced contamination from volume conduction or magnetic field spread. PLoS One, 8(12), 2013.

[11] M. Dumpelmann, T. Ball, and A. Schulze-bonhage. sLORETA allows reliable distributed source reconstruction based on subdural strip and grid Recordings. Human, 33:1172–1188, 2012.

[12] M. Dumpelmann, J. Fell, J. Wellmer, H. Urbach, and C. E. Elger. 3D source localization derived from subdural strip and grid electrodes: a simulation study. Clin. Neurophysiol., 120(6):1061–9, jun 2009.

[13] Q. Fang and D.A. Boas. Tetrahedral mesh generation from volumetric binary and gray-scale images. (Isbi), 2009.

[14] M. D. Fox, D. Zhang, A. Z. Snyder, and M. E. Raichle. The Global Signal and Observed Anticorrelated Resting State Brain Networks. J. Neurophysiol., 101(6):3270–3283, 2009.

[15] P. Fries. A mechanism for cognitive dynamics: neuronal communication through neuronal coherence. Trends Cogn. Sci., 9(10):474–80, oct 2005.

[16] P. Fries. Rhythms for Cognition: Communication through Coherence. Neuron, 88(1):220–235, 2015.

[17] M. Fuchs, M. Wagner, and J. Kastner. Development of volume conductor and source models to localize epileptic Foci. J. Clin. Neurophysiol., 24(2):101–119, 2007.

[18] A. Gramfort, T. Papadopoulo, E. Olivi, and M. Clerc. OpenMEEG : opensource software for quasistatic bioelectromagnetics. pages 1–20, 2010.

[19] R. Grech, T. Cassar, J. Muscat, K. P. Camilleri, S. G. Fabri, M. Zervakis, P. Xanthopoulos, V. Sakkalis, and B.k Vanrumste. Review on solving the inverse problem in EEG source analysis. J. Neuroeng. Rehabil., 5(25), jan 2008.

[20] D.A. Gusnard and M.E. Raichle. Searching for a baseline: Functional imaging and the resting human brain. Nat. Rev. Neurosci., 2(10):685–694, 2001.

[21] C.D. Hacker, A.Z. Snyder, M. Pahwa, M. Corbetta, and E.C. Leuthardt. Frequency-specific electrophysiologic correlates of resting-state fMRI networks. Neuroimage, 149(January):446–457, 2017.

[22] M. Hamalainen, R. Hari, J. Ilmoniemi, J. Knuutila, and O.V. Lounasmaa. Magnetoencephalography≈theory, instrumentation, and applications to noninvasive studies of the working human brain. Rev. Mod. Phys., 65(2), 1993.

[23] P.C. Hansen and D.P. O’Leary. The use of the L-curve in the regularization of discrete ill-posed problems. SIAM J. Sci. Comput., 14(6):1487–1503, 1993.

[24] R. Hari and R. Salmelin. Human cortical oscillation: a neuromagnetic view through the skull. Trends Neurosci., 20(1):44–49, 1997.

[25] B.J. He, A.Z. Snyder, J.M. Zempel, M.D. Smyth, and M.E. Raichle. Electrophysiological correlates of the brain’s intrinsic large-scale functional architecture. Proc Natl Acad Sci USA, 105(41):16039–16044, 2008.

[26] A. Hillebrand, G.R. Barnes, J.L. Bosboom, H.W. Berendse, and C.J. Stam. Frequency-dependent functional connectivity within resting-state networks: An atlas-based MEG beamformer solution. Neuroimage, 59(4):3909–3921, 2012.

[27] R. Hindriks, X.D. Arsiwalla, T. Panagiotaropoulos, M. Besserve, P.F.M.J. Verschure, N.K. Logothetis, and G. Deco. Discrepancies between Multi-Electrode LFP and CSD Phase-Patterns: A Forward Modeling Study. Front. Neural Circuits, 10(July):51, 2016.

[28] J.F. Hipp, D.J. Hawellek, M. Corbetta, M. Siegel, and A.K. Engel. Large-scale cortical correlation structure of spontaneous oscillatory activity. Nat. Neurosci., 15(6):884–90, jun 2012.

[29] J.F. Hipp and M. Siegel. BOLD fMRI correlation reflects frequency-specific neuronal correlation. Curr. Biol., 25(10):1368–1374, 2015.

[30] M. X. Huang, C. W. Huang, A. Robb, A. M. Angeles, S. L. Nichols, D. G. Baker, T. Song, D. L. Harrington, R. J. Theilmann, R. Srinivasan, D. Heister, M. Diwakar, J. M. Canive, J. C. Edgar, Y. H. Chen, Z. Ji, M. Shen, F. El-Gabalawy, M. Levy, R. McLay, J. Webb-Murphy, T. T. Liu, A. Drake, and R. R. Lee. MEG source imaging method using fast L1 minimum-norm and its applications to signals with brain noise and human resting-state source amplitude images. Neuroimage, 84:585–604, 2014.

[31] O Jensen, P Goel, N Kopell, M Pohja, R Hari, and B Ermentrout. On the human sensorimotor-cortex beta rhythm: sources and modeling. Neuroimage, 26(2):347–55, jun 2005.

[32] Y. Jonmohamadi and R.D. Jones. Source-space ICA for MEG source imaging. J Neural Eng, 13(1):16005, 2016.

[33] J. S. Kim, C. H. Im, Y. J. Jung, E. Y. Kim, S. K. Lee, and C. K. Chung. Localization and propagation analysis of ictal source rhythm by electrocorticography. Neuroimage, 52(4):1279–88, oct 2010.

[34] Q. Liu, S. Farahibozorg, and C. Porcaro. Detecting large-scale networks in the human brain using high-density electroencephalography. BioRxiv, pages 1–31, 2016.

[35] L. Manola, B. H. Roelofsen, J. Holsheimer, E. Marani, and J. Geelen. Modelling motor cortex stimulation for chronic pain control: electrical potential field, activating functions and responses of simple nerve fibre models. Med. Biol. Eng. Comput., 43(3):335–343, 2005.

[36] D. Mantini, M. Corbetta, G. L. Romani, G. A. Orban, and W. Vanduffel. Evolutionarily novel functional networks in the human brain? J. Neurosci., 33(8):3259–75, feb 2013.

[37] D. Mantini, S. D. Penna, L. Marzetti, F. de Pasquale, V. Pizzella, M. Corbetta, and G. L. Romani. A Signal-Processing Pipeline for Magnetoencephalography Resting-State Networks. Brain Connect., 1(1):49–59, 2011.

[38] D. Mantini, M.G. Perrucci, C. Del Gratta, G.L. Romani, and M. Corbetta. Electrophysiological signatures of resting state networks in the human brain. Proc. Natl. Acad. Sci., 104(32):13170, 2007.

[39] D.G. Mclaren, K.J. Kosmatka, T.R. Oakes, D. Christopher, S.G. Kohama, J.A. Matochik, D.K. Ingram, and C. Sterling. A population-average MRI-based atlas collection of the rhesus macaque. Neuroimage, 45(1):52–59, 2010.

[40] G. Nolte, O. Bai, L. Wheaton, Z. Mari, S. Vorbach, and M. Hallett. Identifying true brain interaction from EEG data using the imaginary part of coherency. Clin. Neurophysiol., 115(10):2292–307, oct 2004.

[41] Paul L. Nunez and R. Srinivasan. Electric fields of the brain., volume second edi.

[42] G.C. O’Neill, E.L. Barratt, B.A.E. Hunt, P.K. Tewarie, and M.J. Brookes. Measuring electrophysiological connectivity by power envelope correlation: a technical review on MEG methods. Phys. Med. Biol., 60(21):R271–R295, 2015.

[43] T.F. Oostendorp and A. van Oosterom. Source parameter estimation in inhomogeneous volume conductors of arbitrary shape. IEEE Trans. Biomed. Eng., 36(3):382–391, 1989.

[44] R. Oostenveld, P. Fries, E. Maris, and J.-M. Schoffelen. FieldTrip: Open source software for advanced analysis of MEG, EEG, and invasive electrophysiological data. Comput. Intell. Neurosci., page 156869, jan 2011.

[45] E. Pereda, R. Quian, and J. Bhattacharya. Nonlinear multivariate analysis of neurophysiological signals. Prog. Neurobiol., 77:1–37, 2005.

[46] Y. Petrov. Harmony: EEG/MEG linear inverse source reconstruction in the anatomical basis of spherical harmonics. PLoS One, 7(10):e44439, jan 2012.

[47] G. Ramantani, D. Cosandier-Rimele, A. Schulze-Bonhage, L. Maillard, J. Zentner, and M. Diimpelmann. Source reconstruction based on subdural EEG recordings adds to the presurgical evaluation in refractory frontal lobe epilepsy. Clin. Neurophysiol., 124(3):481–91, mar 2013.

[48] P. Ritter, M. Moosmann, and A. Villringer. Rolandic alpha and beta EEG rhythms’ strengths are inversely related to fMRI-BOLD signal in primary somatosensory and motor cortex. Hum. Bram Mapp., 30(4):1168–1187, 2009.

[49] B. Rubehn, C. Bosman, R. Oostenveld, P. Fries, and T. Stieglitz. A MEMS-based flexible multichannel ECoG-electrode array. J. Neural Eng., 6(3):036003, jun 2009.

[50] D. Rubino, K.A. Robbins, and N.G. Hatsopoulos. Propagating waves mediate information transfer in the motor cortex. Nat. Neurosci., 9(12):1549–57, dec 2006.

[51] J.N. Sanes and J.P. Donoghue. Oscillations in local field potentials of the primate motor cortex during voluntary movement. Proc. Natl. Acad. Sci. U. S. A., 90(10):4470–4474, 1993.

[52] Marcus Siems, Anna Antonia Pape, Joerg F. Hipp, and Markus Siegel. Measuring the cortical correlation structure of spontaneous oscillatory activity with EEG and MEG. Neuroimage, 129:345–355, 2016.

[53] C.J. Stam, G. Nolte, and A. Daffertshofer. Phase lag index: assessment of functional connectivity from multi channel EEG and MEG with diminished bias from common sources. Hum. Brain Mapp., 28(11):1178–93, nov 2007.

[54] K. Takahashi, S. Kim, T.P. Coleman, K.A. Brown, A.J. Suminski, M.D. Best, and N.G. Hatsopoulos. Large-scale spatiotemporal spike patterning consistent with wave propagation in motor cortex. Nat. Commun., 6(May):7169, jan 2015.

[55] D. C. Van Essen, H. A. Drury, J. Dickson, J. Harwell, D. Hanlon, and C. H. Anderson. An Integrated Software Suite for Surface-based Analyses of Cerebral Cortex. 8(5):443–459, 2001.

[56] Y. Zhang, W. Drongelen, van, M. Kohrman, and B. He. Neuroimage Three-dimensional brain current source reconstruction from intra-cranial ECoG recordings. Neuroimage, 42:683–695, 2008.

